# Sequential Switching Through IgG1 is Redundant for Allergic Reactivity and Memory to Allergens

**DOI:** 10.1101/2023.05.27.542563

**Authors:** Joshua F.E. Koenig, Adam Wade-Vallance, Rodrigo Jiménez-Saiz, Kelly Bruton, Siyon Gadkar, Emily Grydziuszko, Tina Walker, Melissa E. Gordon, Susan Waserman, Manel Jordana

## Abstract

Allergic reactions to foods are driven by allergen-binding immunoglobulin (Ig)E antibodies. IgE-expressing cells can be generated through a sequential class switching pathway where activated B cells first switch to an intermediary isotype, most frequently IgG1, and then to IgE. It has been proposed that sequential class switch recombination is important in generating high affinity IgE, augmenting anaphylactic reactions, and in holding the memory of IgE responses. Here, we observed surprising redundancy of sequential switching through IgG1 for the functional affinity of the IgE repertoire against multiple food allergens as well as for the ability of IgE to elicit anaphylaxis. We further found that sequential switching via IgG1 was irrelevant for allergic memory. These results indicate that allergen-specific IgG1 B cells are redundant in sensitization, anaphylaxis, and food allergy persistence, thereby implicating other switching pathways as important considerations in the development of therapeutics for allergic diseases.

## Introduction

Food allergy is an immunological disorder associated with a significant health, quality of life, and financial burden^1, 2^. The production and persistence of allergen-specific IgE determines allergic reactivity. Therefore, understanding the requirements for IgE production of sufficient binding capacity to trigger anaphylaxis is critical for the development of preventative and curative treatments in food allergy.

Studies in mice have revealed that IgE-expressing B cells are largely excluded from the germinal centre (GC), which is the classical pathway for B cells to attain high affinity and longevity^3–6^. B cells augment their affinity for antigen in the GC through iterative rounds of *Ig* gene mutation and positive selection. These positively selected GC B cells are a critical source of long-lived cells, including memory B cells (MBCs) and plasma cells. Early work which imaged IgE-expressing cells in mice argued that IgE B cells are absent from the GC^5^. Later work revealed that IgE B cells constitute a rare fraction of the early GC but are rapidly eliminated as the reaction progresses^3, 7^. Congruent with these findings, IgE B cells are severely or even completely constrained in MBC and long-lived plasma cell differentiation^3, 8^. These observations contrast with lifelong reactivity in most patients allergic to peanuts, tree nuts, fish and shellfish, as well as with the detection of high affinity IgE clones in some allergic patients^9^.

A suggested resolution for these discordant observations is the sequential switching hypothesis, which argues that IgE-secreting cells emerge from B cells that previously underwent class switch recombination (CSR) to a non-IgE isotype, usually IgG1. This hypothesis originated from early work which found that remnants of the IgG1 switch region were present in the genomic switch regions of some IgE-expressing B cells stimulated *in vitro* or from parasitized mice^10, 11^. This pathway purportedly overcomes the aforementioned limitations of IgE B cells: IgG1-expressing B cells can gain high affinity mutations in the GC prior to sequential CSR to IgE, and long lived IgG1 MBCs can undergo sequential CSR to IgE upon allergen re-exposure to maintain IgE _titres_^5, 12, 13^.

Evidence supporting a role for sequential CSR through IgG1 in IgE production is conflicting. In some instances, the production of IgE has been noted to lag behind the peak production of IgG1^5, 11, 14^, perhaps indicative of the additional time required for sequential CSR from IgG1 to IgE. However, mice that are unable to undergo CSR to IgG1 retained similar levels of IgE relative to wildtype (WT) mice^15^. Due to the normal magnitude of IgE responses in the absence of IgG1 CSR, it was suggested that sequential CSR through IgG1 is an obligatory step for the generation of high-affinity IgE, rather than for IgE *per se*^5^. In a hapten-based immunization system, IgG1-deficient (IgG1-def) mice had normal levels of specific IgE but with impaired binding affinity, supporting this contention^16^. In contrast, epitope scanning of human peanut-allergic sera from patients avoiding foods found incomplete overlap of the IgG and IgE repertoire^17^; many epitopes were bound by IgE but not IgG, suggesting that a cellular IgG intermediate may not naturally exist for those IgE-secreting cells. However, the overlap of the compartment appears to increase following recall allergen exposure through immunotherapy^17, 18^, suggesting that recall IgE can be derived from IgG MBCs. This is consistent with mouse models which have demonstrated that IgG1 MBCs can undergo sequential CSR to IgE upon allergen re-exposure^5, 13^. Ultimately, whether sequential CSR through IgG1 is required for clinical reactivity or memory of IgE responses against allergens remains to be elucidated.

To address this gap, we compared the allergic reactivity of IgG1-def and -sufficient mice in well-established mouse models of food allergy. We found that IgG1-def mice had similar levels and polyclonal affinity of allergen-specific IgE, and similar severity of anaphylaxis upon challenge with either egg or peanut allergens. However, IgG1-def mice produced lower affinity anti-hapten IgE and exhibited reduced clinical reactivity upon challenge with hapten conjugates, suggesting that different types of antigens have different CSR requirements. Furthermore, IgG1-def and wildtype (WT) mice mounted similarly robust recall IgE responses at memory timepoints that drove severe anaphylaxis upon challenge, indicating that features of MBCs other than isotype define their capacity to hold IgE memory. Therefore, sequential CSR through IgG1 is redundant to generate anaphylactic IgE and to hold IgE memory.

## Results and Discussion

### Sequential switching through IgG1 is redundant for IgE-mediated reactivity to food allergens

To evaluate the contribution of sequential CSR through IgG1 to allergic responses to food allergens, we utilized hMT mice (hereafter referred to as IgG1-def) which lack the IL-4 sensitive elements of the Iγ1 promoter region, thereby preventing CSR to IgG1^15^. We sensitized WT and IgG1-def mice to a common egg allergen (ovalbumin, OVA), or to peanut (PN), which contains several allergens, using a well-characterized intragastric model of food allergy^19, 20^. This regimen yields allergen-specific IgG1 and IgE which mediate anaphylactic reactions upon allergen challenge. In mice, anaphylaxis is measured by a drop in core body temperature, increased hematocrit (indicative of vascular leakage), and clinical signs ranging from inner ear scratching to seizure (see Materials and Methods)^19, 20^. IgG1-def mice had an expected absence of serum allergen-specific IgG1 following sensitization with either OVA or PN (Fig. S1A). IgG1-def mice also generated fewer allergen-specific Igs (Fig. S1B), indicating that the absence of IgG1 was not fully compensated by other isotypes. However, the absence of sequential CSR through IgG1 did not impact the magnitude of allergen-specific IgE production (Fig. 1A), consistent with previous reports^15, 16^. Upon allergen challenge, IgG1-def and WT mice experienced a similar drop in core body temperature (Fig. 1B), increase in hematocrit (Fig. 1C), and clinical signs (Fig. S1C). Altogether, these results demonstrate that sequential CSR through IgG1 is not required for clinical reactivity against food allergens.

**Figure 1:**
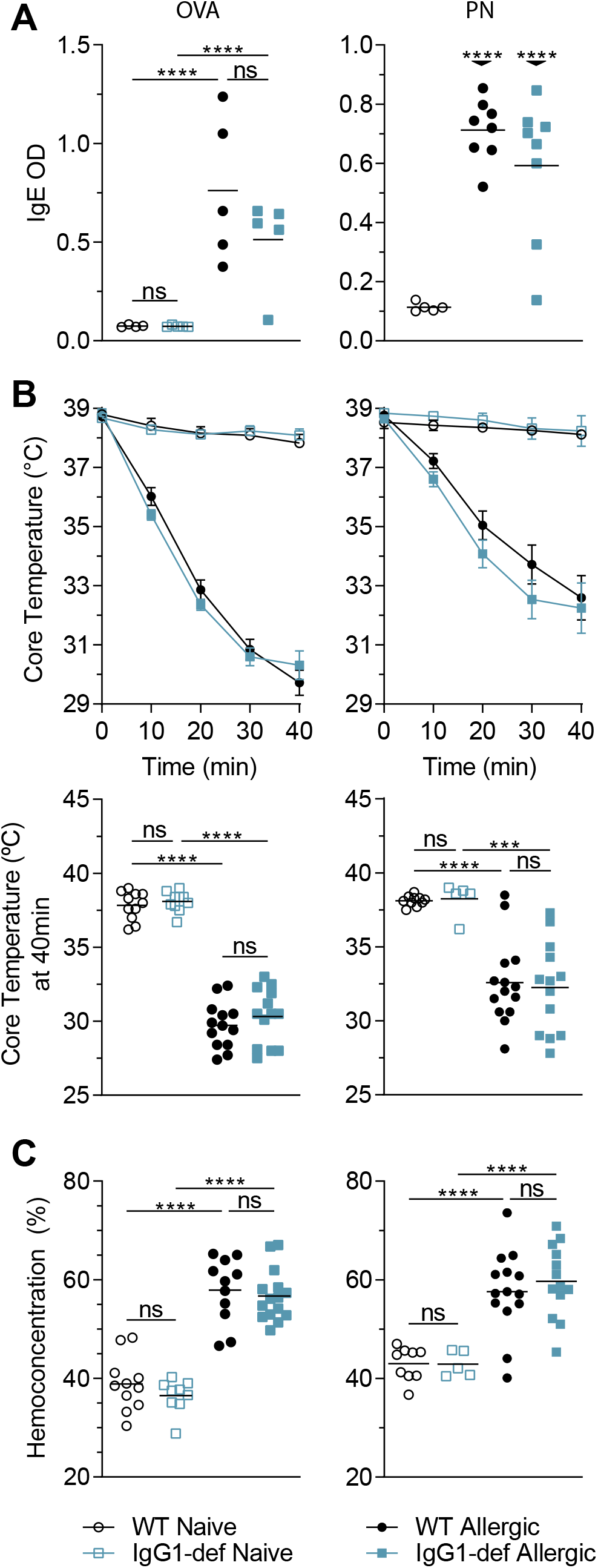
Sensitized IgG1-deficient and WT mice have similar allergic reactivity to foods. (A-C) Wildtype (WT) and IgG1-deficient (IgG1-def) mice were sensitized to either OVA (left column) or PN (right column) by 4 intragastric exposures of allergen + cholera toxin, as described in Materials and Methods, or were untreated (naïve). **(A)** Quantification of serum allergen-specific IgE in sera by ELISA at 450nm. **(B-C)** Two weeks following sensitization mice were challenged by intraperitoneal injection of food allergen. **(B)** Depicted are measurements of core temperature over 40 minutes following challenge by rectal probe (top) with statistical analysis (bottom) performed at 40 minutes post-challenge. **(C)** Quantification of hematocrit in blood collected at 40 minutes post-challenge. ***, P < 0.001, ****, P < 0.0001 (one way ANOVA with Tukey’s post- test comparing the mean of each group with the mean of every other group [A-C]). Statistical notations accompanied by downward arrows (A, right) indicate comparisons between the indicated group and the naïve control group, whereas notations atop bars indicate comparisons between the bridged groups. Dots represent group means (B, top panels), readings from individual mice (B, bottom panels), or measurements of samples derived from individual mice (A, C). Bars within groups represent the mean. Data are representative (A) or pooled from (B-C) three independent experiments.

The classical pathway of anaphylaxis involves the allergen-mediated cross-linking of IgE bound by FcεRI on the surface of mast cells and basophils, resulting in the release of allergic mediators. However, an “alternative pathway” driven by IgG-allergen immune complexes, which ligate FcγRII/III on the surface of macrophages, can also contribute to anaphylaxis^21^. To confirm that the anaphylaxis experienced by IgG1-def mice was IgE-mediated, we blocked the alternative pathway by the well-established approach of administering an anti-FcγRII/III antibody prior to challenge^21, 22^. FcγRII/III-blockade issued a 50% reduction in the level of hypothermia in allergic WT mice following allergen challenge (Fig. S2A), consistent with our previous report^21^. We did not detect a difference in hemoconcentration (Fig. S2B), though FcγRII/III-blockade did partially protect WT mice against clinical symptoms of anaphylaxis relative to isotype control-treated animals (Fig. S2C). Conversely, IgG1-def mice were not significantly protected from anaphylaxis when treated with anti-FcγRII/III (Fig. S2A-C), indicating that they underwent IgE-mediated anaphylaxis. Therefore, sequential CSR through IgG1 is redundant for IgE-mediated anaphylaxis against food allergens.

### Sequential switching through IgG1 contributes to IgE-mediated reactivity to haptens

Our observation that IgG1-def mice generated anaphylactic IgE antibodies appears to contrast with a study reporting that sequential CSR through IgG1 was required for high affinity IgE production upon systemic sensitization with the hapten 4-hydroxy-3-nitrophenyl (NP)^16^. We hypothesized that the different routes of sensitization used (systemic *vs*. intragastric) might explain these discordant results, and so we sensitized mice intragastrically with NP conjugated to OVA (NP-OVA). To measure allergic reactivity against NP rather than OVA, we challenged mice with NP conjugated to bovine serum albumin (NP-BSA). Consistent with our previous experiments, IgG1- def mice lacked NP-specific IgG1, had reduced total NP-specific Igs, and had similar production of NP-specific IgE relative to WT animals (Fig. S3A-C). We compared the polyclonal binding affinity of NP-IgE between WT and IgG1-def mice by calculating the ratio of serum IgE binding to lowly- *vs.* highly-conjugated NP-BSA^23^. IgE affinity was significantly reduced in hapten- sensitized IgG1-def mice compared to WT, as had been previously described in the systemic immunization model (Fig. S3C-D). Concordantly, IgG1-def mice were partially protected from hypothermia and vascular leakage and exhibited less severe clinical signs compared to WT mice (Fig. S3E-G). Together with our earlier results, these findings reveal a fundamental difference in the contribution of sequential CSR through IgG1 to IgE-mediated allergic reactivity to haptens compared to proteins, independent of sensitization route.

### Sequential switching through IgG1 is redundant for the functional polyclonal affinity of food allergen-specific IgE

Given the divergent importance of sequential CSR through IgG1 in hapten and food allergen systems, we next asked whether IgE affinity was impaired against food allergens in the absence of sequential CSR through IgG1. Firstly, we reasoned that potent clinical reactivity driven by high affinity antibodies would be required in the context of limiting allergen challenge doses. Indeed, adjusting the challenge dose from 0.25 mg (Fig. 2A) to 2.5 mg (Fig. 1B) resulted in dose-dependent changes in the reactivity of WT mice. Across all challenge doses, IgG1-def mice experienced equivalent or greater hypothermia (Fig. 2A-B) and hematocrit (Fig. 2C) compared to WT mice, suggestive of equivalent or greater polyclonal IgE affinity. However, the greater reactivity in IgG1- def mice could also reflect their reduced total allergen-specific Ig (Fig. S1B) and a corresponding weaker blocking of allergen-IgE interactions on mast cells and basophils. To measure IgE function in an isolated manner, we evaluated the capacity of IgE from IgG1-def and WT mice to induce degranulation *in vitro* of rat basophil leukemia-2H3 (RBL) cells, a common model for mouse mast cells^24^. RBL cells were incubated with serum from sensitized IgG1-def or WT mice, washed to remove unbound antibody, and then challenged with allergen. Serum from IgG1-def and WT mice equivalently primed RBL cells for degranulation (Fig. 2D). To confirm that RBL degranulation was IgE-mediated, we heat-inactivated the serum at 56°C prior to incubation with the RBL cells. Unlike other Igs, IgE is irreversibly denatured at 56°C due to a heat-labile domain which is not present in other antibody isotypes^25, 26^. Following heat-inactivation, RBL cells did not degranulate when sensitized with either IgG1-def or WT serum, confirming that the reactivity seen in this system was IgE-mediated (Fig. 2D). Altogether, these results support that IgG1-def and WT mice produced IgE of similar polyclonal affinity.

**Figure 2:**
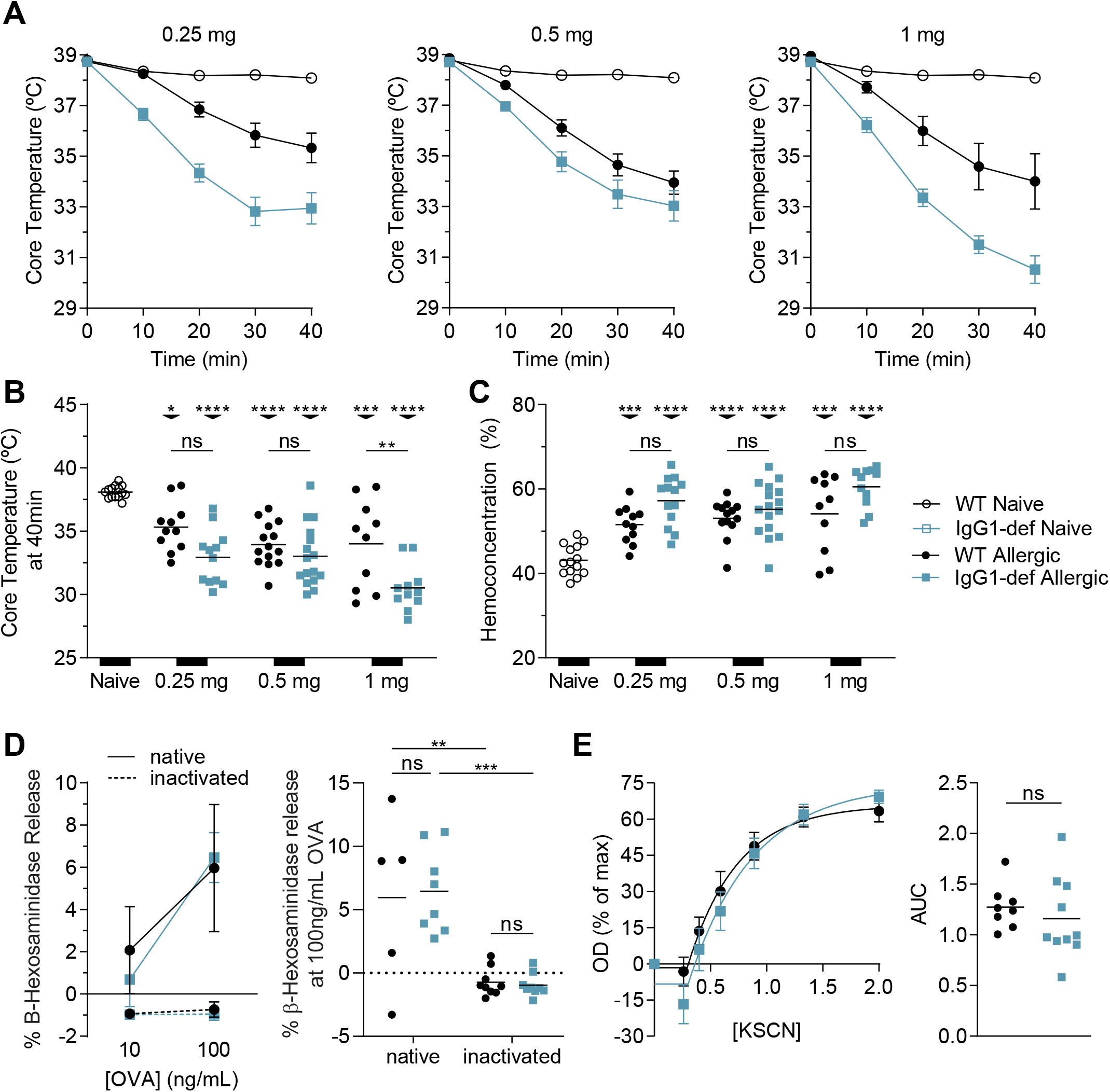
IgG1-def and WT mice generate allergen-specific IgE of similar functional affinity. (A-E) WT and IgG1-def mice were sensitized to either PN or OVA or were left naïve. **(A)** Measurement of core body temperature change over time by rectal probe following PN challenge at one of three doses: 0.25mg (left), 0.5mg (middle) and 1mg (right). **(B)** Statistical analysis of data presented in Panel A at the 40 minutes post-challenge. **(C)** Quantification of hemoconcentration in blood collected at 40 minutes post-challenge. **(D)** β-hexosaminidase release from RBL-2H3 cells challenged with OVA after incubation with serum from sensitized mice. Statistical analysis for the 100ng/mL dose is shown in the right plot. **(E)** Average curve of the % reduction in OVA-IgE ELISA signal as a function of increasing KSCN concentration (left), and quantification of the area under the curve (AUC) for individual samples (right). *, P<0.05; **, P<0.01; ***, P < 0.001; ****, P < 0.0001 (one way ANOVA with Tukey’s post-test comparing the mean of each group with the mean of every other group [A-C]). Dots represent the group mean (A, left panels of D and E), readings from individual mice (B) or measurements of samples from individual mice (C, right panels of D and E). Statistical notations accompanied by downward arrows indicate comparisons between the indicated group and the naïve control group, whereas notations atop bars indicate comparisons between the bridged groups. Bars within groups represent the mean. Data are pooled from (A-C) or representative of (D-E) three independent experiments.

To strengthen this conclusion, we directly compared the allergen-binding capacity of allergen-specific IgE from IgG1-def mice and WT mice using an affinity-dependent ELISA technique. Briefly, we adapted our allergen-specific IgE ELISA to include an elution step using the chaotrope thiocyanate which dose-dependently interrupts the allergen-IgE interaction. High-affinity antibodies are resistant to elution, allowing polyclonal binding affinity to be evaluated using elution curves^27^. Allergen-specific IgE from IgG1-def and WT mice eluted at similar concentrations of thiocyanate (Fig. 2E), indicating that they had similar polyclonal binding affinity for allergen. Cumulatively, these data indicate that IgE produced against food allergens in the absence of sequential CSR through IgG1 has intact functional polyclonal affinity for allergen.

### Compensation between isotypes in the germinal center

Given its central role in the production of affinity-matured B cells, we reasoned that alterations in the GC of IgG1-def mice could explain their intact IgE affinity and reactivity against food allergens. To investigate the allergen-specific GC response, we detected OVA-specific GC B cells (defined as CD19^+^GL7^+^CD95^+^) at the previously reported peak of the primary response^28^ in the mesenteric lymph nodes using monomeric OVA-FITC (Fig. 3A). IgG1-def mice had a lower proportion and number of OVA^+^ GC B cells compared to WT mice (Fig. 3B), indicating an inability to fully compensate for the absence of the IgG1 GC B cells, which constitute ∼90% of the antigen-specific GC B cell compartment in WT mice (Fig. 3C). While the overall GC response of IgG1-def mice was reduced, they exhibited an increased frequency and absolute number of OVA-specific IgE-, IgM- and IgG3-expressing GC B cells compared to WT mice (Fig 3D-F). They also had comparable or slightly elevated number and frequency of OVA-specific IgG2b- and IgG2c-expressing GC B cells (Fig. 3G,H). The compensatory numerical increases in non-IgG1 GC B cells, especially IgE, IgM, and IgG3, in IgG1-def mice provide a cellular basis for their intact IgE affinity and allergic reactivity.

**Figure 3:**
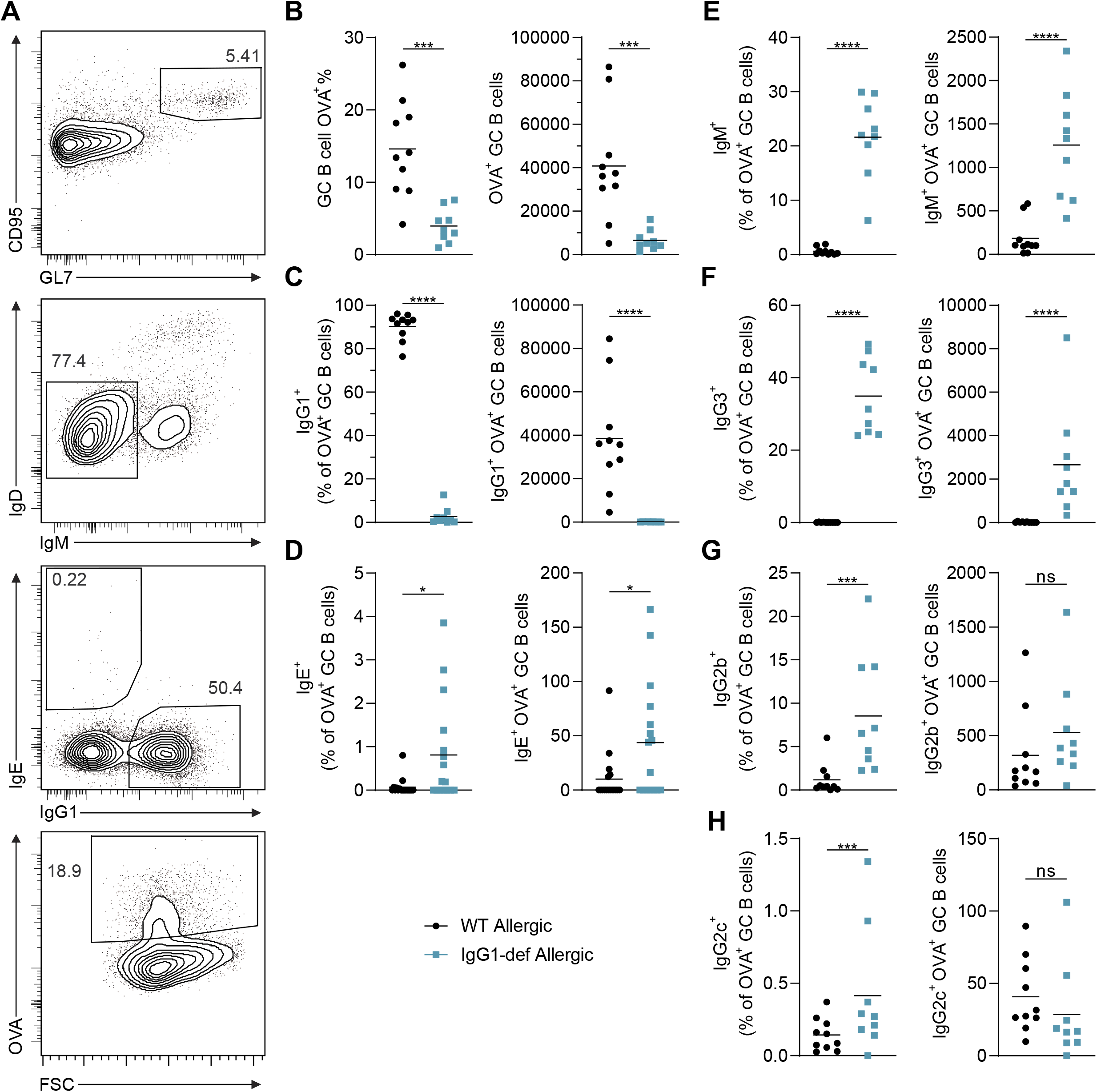
Isotype compensation in the IgG1-def GC. (A-I) IgG1-def and WT mice were sensitized to OVA and the GC response in the mesenteric lymph nodes were assessed by flow cytometry 6 d later. **(A)** Representative contour plots depicting the hierarchical gating strategy used to identify allergen-specific GC B cells. Cells were pre-gated as CD19^+^ live singlets, then as CD95^+^ GL7^+^ GC B cells, IgM^-^IgD^-^, by isotype (shown for IgE and IgG1), and as allergen binding using a monomeric OVA-FITC probe. **(B-H)** The proportion (left) and absolute counts (right) of total allergen-specific GC B cells (B) or of allergen-specific GC B cells of the indicated isotype (C-H). *, p<0.05; ***, P < 0.001; ****, P < 0.0001 (unpaired t test). Dots represent samples from individual mice and bars within groups represent the mean. Data are representative of (A) or pooled from (B-H) two (E-H) or three (B-D) independent experiments.

### Sequential switching through IgG1 is redundant for memory IgE responses

Given the importance of the GC in the production of MBCs and that IgG1 MBCs are thought to be the primary reservoir of IgE memory in both mice and humans,^12, 13, 29^ we next sought to determine the importance of IgG1 to the IgE recall response and, by extension, allergic memory. To investigate this, we sensitized IgG1-def and WT mice and waited until clinical reactivity waned at 10 months post-sensitization (Fig. 4A). We have previously reported that oral exposure to allergen without adjuvant at this timepoint results in a robust secondary response that replenishes allergen-specific IgE titres and allergic reactivity^28^. Titres of allergen-specific IgE in IgG1-def mice were no longer detectable above background at 20 weeks post-sensitization, while most WT mice had detectable allergen-specific IgE until 45 weeks post-sensitization (Fig. 4B). Consistent with our previous report, WT and IgG1-def mice lost the majority of their clinical reactivity after 10 months (Fig. 4C). Upon re-exposure, sensitized IgG1-def and WT mice exhibited a similar regeneration of allergen-specific IgE (Fig. 4D). Challenging re-exposed allergic mice with allergen revealed comparable hypothermia (Fig. 4C), hematocrit (Fig. 4E), and clinical signs (Fig. 4F) between IgG1-def and WT mice. These data demonstrate that sequential CSR through IgG1 is not a critical pathway for IgE recall responses or allergic memory.

**Figure 4:**
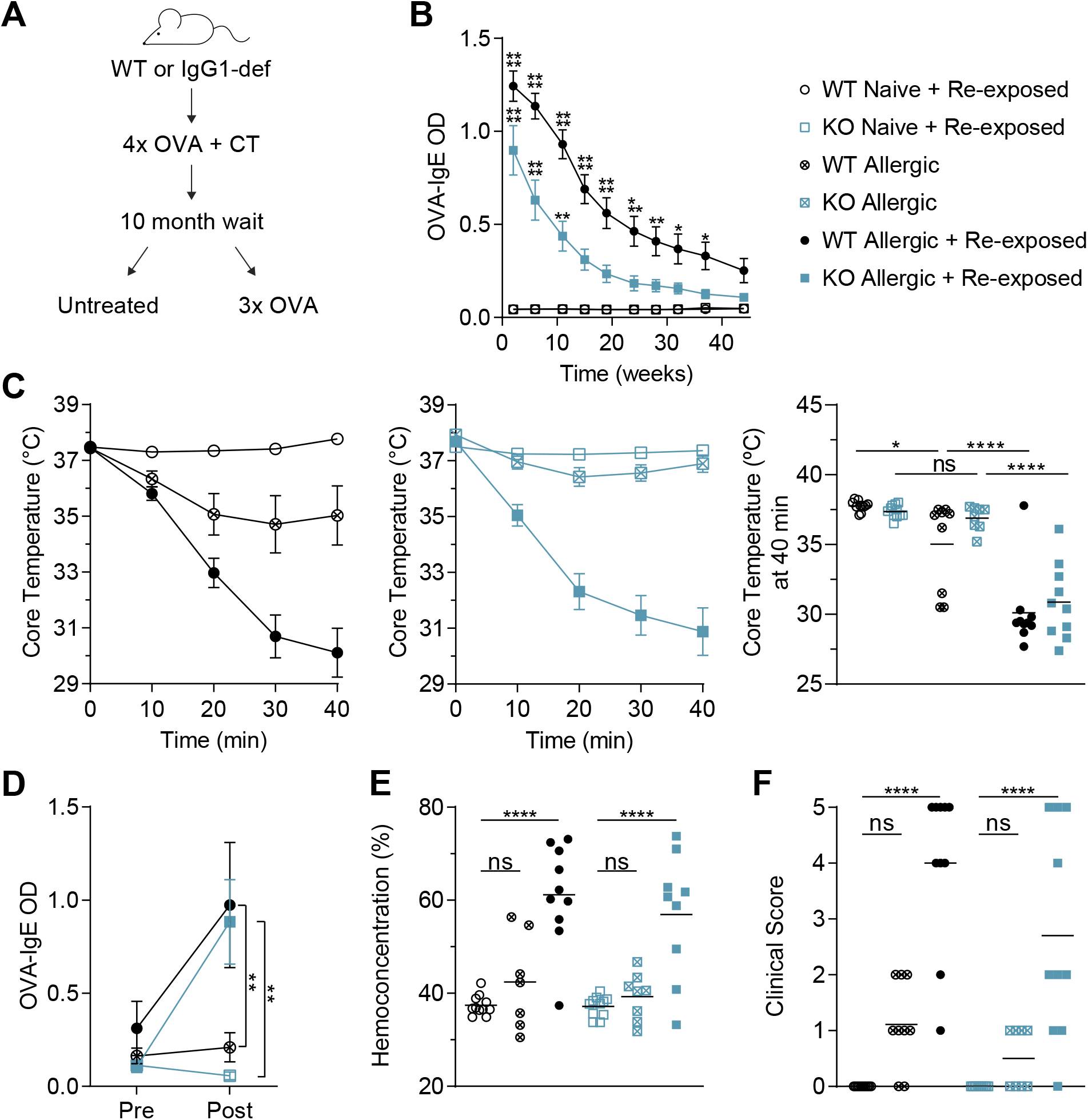
IgG1-def and WT mice have equivalent allergic memory. (A) Schematic depicting the allergic memory model used in panels B-F. Mice were left naïve (hollow symbols) or sensitized with OVA + cholera toxin. Following sensitization, allergic reactivity was allowed to wane over a 10-month period, after which one group of allergic mice was given three oral re-exposures to allergen prior to challenge (solid symbols) whereas the other was not (hollow symbols with cross. **(B)** Quantification of OVA-specific IgE *vs.* weeks following sensitization. Statistical notations indicate comparisons between the below allergic mice and the naïve controls of the same genotype. When notations are not provided the comparisons are ns. **(C)** Core temperature change over time following challenge of WT (left) and IgG1-def (centre) mice with accompanying statistical analysis (right). **(D)** Serum OVA-specific IgE measured by ELISA at 450nm before (Pre) and after (Post) the oral re-exposures at 10 months post-sensitization. Statistical notations indicate comparisons between the bridged groups at the “Post” timepoint. **(E)** Hematocrit in blood collected at 40 minutes post-challenge. **(F)** Clinical signs of anaphylaxis assessed over 40 minutes following allergen challenge. *, p<0.05; **, P<0.01; ***, P < 0.001; ****, P < 0.0001 (two-way ANOVA comparing the mean of each group with the mean of every other group [A], one way ANOVA comparing the mean of each group with the mean of every other group [C, right]), one way ANOVA comparing the mean of the indicated pairs of groups [D, E, F]. Dots represent the group mean (B, C left and centre panels, D), readings from individual mice (C right panel, F) or measurements of samples from individual mice (E). (C-F) Statistical notations atop bars indicate comparisons between the bridged groups. Bars within groups represent the mean. Data are representative of (B, D) or pooled from (C, E-F) two independent experiments.

In summary, we have shown that IgG1-def mice can undergo severe, IgE-mediated allergic reactions against food allergens. Whereas sequential CSR through IgG1 contributes to allergic reactivity and IgE affinity to haptens, it plays no mandatory role in allergic reactivity or IgE affinity to food allergens. At the cellular level, the absence of IgG1 B cells alters GC composition, most notably increasing IgE, IgM, and IgG3 GC B cells, indicating that affinity maturation can occur in IgG1-def mice. We also demonstrated that allergic recall responses to food allergens were intact and equivalent in IgG1-def mice relative to WT controls. Overall, our findings indicate that sequential CSR through IgG1 is redundant for both of the two core features of allergic responses: the production of anaphylactic IgE and allergic memory. These results call for greater scrutiny of the prevailing notions that sequential CSR and high-affinity IgE are critical for anaphylactic reactions and have important implications for the development of allergy therapeutics.

Our observation that IgG1 is redundant for IgE-mediated reactivity to food allergens appears to contradict classical associations linking IgE and IgG1 responses. One possible explanation is that IgE responses are less dependent on IgG1 than previously appreciated. The original observations that formed the basis for the IgG1 sequential CSR hypothesis were the shared induction of IgG1 and IgE by IL-4 and the frequent identification of γ1 switch remnants in IgE B cells^11, 30, 31^. However, we have since learned more about mechanisms which uniquely restrict IgE and not IgG1 responses (*e.g.* IL-21)^7, 32, 33^, exposing that IgE and IgG1 may be less tightly linked than once thought. Furthermore, it is generally assumed that the rate at which switch remnants are identified represents the minimal possible contribution of IgG1 intermediaries given that only a portion of switching events leave these remnants^31^. However, switch remnants can be present on the productively recombined IgH allele or on the non-recombined inactive allele^34^, and many published analyses do not discriminate between the two. Further, the recent discovery of a ‘reverse sequential switching pathway’ in which the IgG1 and IgE loci recombine first, prior to recombination with the IgM locus, suggests that some reported switch remnants are not indicative of a cellular IgG1-expressing intermediate. Therefore, switch remnant-based analyses could also over-estimate IgG1 B cell ancestry, and thus do not provide concrete evidence for understanding the CSR history of a cell.

A second possible explanation for prior associations of IgG1 with IgE is that most allergen-specific B cells switch to IgG1 (>90% of OVA-specific GC B cells in the present study; Fig. 3C), therefore the most likely cell to undergo CSR to IgE would be an IgG1-expressing cell. By this model, the increased abundance of other isotypes of non-IgE B cells (*e.g.*, IgM, IgG3) in IgG1-def mice fill the niche of IgE intermediaries normally occupied by IgG1, resulting in intact sensitization and memory responses. However, this model cannot explain results from studies which interfered with the IgG1 response rather than eliminating it altogether. For example, IgE production was impaired by treating IL-4- and endotoxin-stimulated B cell cultures with anti-IgG1 antibodies^31^ or swapping the extracellular domains of IgG1 and IgE^35^. Therefore, a more nuanced possibility is that IgG1 is so dominant in the allergen-specific B cell response of WT mice that IgG1-independent pathways for IgE production are suppressed. Only when IgG1 switching is completely blocked can these pathways support full IgE switching and affinity, as in the present study. One potential mechanism for this suppression is the dominance of IgG1 within the GC. Previous work has shown that IgG GC B cells are more competitive than their IgM counterparts, even with the same variable region sequence, indicating that the constant region of the B cell receptor is important to GC selection^36^. Indeed, we observed robust populations of IgM, IgE, and IgG3 GC B cells in IgG1-def mice (Fig. 3E). IgM and IgE GC B cells represent especially strong candidates as intermediates for IgE-secreting cells in IgG1-def mice; according to the text of a prior study (data not depicted), IgG1- def mice showed no evidence of increased sequential switching through non-IgG1 isotypes and therefore the authors concluded that all switching to IgE in IgG1-def mice is direct^16^.

Our findings demonstrate the importance of investigating food allergy using *bona fide* food allergens. This is evident in the differential requirement for CSR through IgG1 for robust allergic reactivity against haptens *vs.* allergens. This difference may be explained by the observation that immunization with haptens induces a GC which purifies for affinity-enhancing mutations, whereas complex antigen-directed GCs maintain epitope diversity at the cost of affinity enhancement^37^. In the setting of PN allergy, the epitope diversity of specific IgE is a stronger determinant of reactivity than polyclonal IgE affinity,^17, 38–42^ findings echoed by a recent mechanistic study of mast cell activation by PN-specific IgE^40^. Hapten-based models may amplify the minor contribution of IgE affinity to allergic reactivity because there is only one epitope to bind, and therefore IgE affinity dictates sensing by mast cells in opposition to blocking Ig. In food allergy, the immune response selects for epitope diversity, which is the predominant determinant of clinical reactivity and does not require IgG1 GC B cells^43^, the GC^28^, or sequential CSR through IgG1 (Fig. 1).

Importantly, we found that the absence of IgG1 had no impact on allergic recall responses. These observations challenge the widespread notion that IgG1 MBCs act as a critical reservoir for secondary IgE responses^12, 13, 18, 44^. While our data could be interpreted to mean that IgG1 MBCs do not contribute to IgE recall responses in the WT setting, clonal connections in humans and adoptive transfer experiments in mice suggest that they likely do contribute. An alternate hypothesis consistent with our findings is that MBC phenotype, rather than isotype, is the primary determinant of its tendency to contribute to IgE recall responses. In support of this, a recently identified type 2 polarized MBC (MBC2) phenotype was found to be the primary clonal relative to IgE plasma cells following chronic allergen exposure in humans^45–47^. While MBC2s are enriched in IgG1- expressing cells, our data demonstrate that IgG1 is not a defining feature of an MBC that later gives rise to IgE-secreting cells^45^. This is consistent with B cell receptor repertoire analysis which found that all upstream isotypes are clonally related to IgE in circulation and along the gastrointestinal tract^48–50^. Overall, our data call for a re-evaluation of the perceived critical role for IgG1 MBCs and sequential CSR through IgG1 in the emergence and persistence of food allergy. Also, our findings indicate that to achieve transformative treatments for IgE-mediated diseases both IgG1- and non-IgG1 MBCs should be considered as therapeutic targets.

## Materials and Methods

### Mice

Age-, sex-, and vendor-matched control mice were used for all experiments. Mice were maintained in biohazard specific pathogen-free conditions on a 12-hour light-dark cycle with low-fat food and water *ad libitum*. Where possible, littermate controls were used. C57Bl/6 mice were purchased from Charles River Laboratories. IgG1-deficient (hMT) mice were provided by Dr. Amy Kenter (University of Illinois). All animal procedures were approved by McMaster University’s Animal Research Ethics Board.

### Intragastric model of sensitization

Mice were sensitized with either 3.75 mg All-Natural Smooth Peanut Butter (Kraft Heinz, Northfield, USA), 1 mg ovalbumin (OVA) (MilliporeSigma, A5378), or 1 mg 4-hydroxy-3- nitrophenyl (NP)-OVA (LGC Biosearch Technologies, N-5051-100) by oral gavage with 5 μg cholera toxin (List Biological Labs, 100B) in 0.5 ml PBS. Gavages were performed once weekly for 4 weeks. Serum was collected from retro-orbital bleeds between 2 weeks and 12 months post-sensitization for serum Ig analyses. Two-weeks following the last gavage, mice were challenged intraperitoneally with 0.25 - 2.5 mg OVA or crude peanut extract (CPE; Stallergenes Greer USA, XPF171D3A25). In NP sensitization experiments, NP-BSA (LGC Biosearch Technologies, N5050H-100) was used for challenge to measure reactivity specific to NP, rather than the OVA- conjugate used for sensitization. Antigens used for sensitization and challenge are noted in the figure legend. Core temperature was measured at 10-minute intervals for 40 minutes post-challenge using a rectal probe (VWR International, 23226-656). Hemoconcentration was measured using the HemataSTAT II centrifugation device (VWR International, 14221-620). Clinical signs were scored by a blinded research technician; scoring criteria are: 0 = no clinical signs, 1 = hind leg scratching in the ear, 2 = reduced movement, 3 = motionless, 4 = no response to whisker stimuli, 5 = moribund, seizure, or death. The NP:carrier ratio of NP-conjugated reagents depended on availability from the supplier; NP-OVA ranged from 16-22 NP moieties/molecule, and NP-BSA ranged from 27-30. To block IgG-mediated anaphylaxis, 500 μg anti-CD16/32 (Clone 2.4G2; Bio X Cell, BE0307) in 500 μl PBS was administered by intraperitoneal injection 24 hours prior to challenge.

For memory experiments, mice were sensitized as above and then left to desensitize over 10 months. Memory responses were induced by re-exposing the mice with 3 gavages of 1 mg OVA alone (no adjuvant), one week apart.

### Antigen-specific Ig ELISA

Blood was collected by retro-orbital bleed and centrifuged at ≥9000 g to separate serum and cellular fractions. Serum was removed and stored at -20℃ for ELISA.

ELISAs were performed in a 96-well flat bottom polystyrene plate (VWR, 4394554). Plates were washed using a Tecan Hydroflex Plate Washer and read using a Thermo Scientific Multiskan FC. No-sample controls were included in all assays. The average optical density of the no-sample control wells was subtracted from all samples prior to plotting. In cases where subtraction would result in negative values for some samples, instead the largest value that would yield an OD of at least 0.01 would be subtracted to ensure compatibility with log transformation (see statistical analysis below).

#### IgG1, IgG3, Total Ig

Plates were coated with either 4 μg/ml OVA, NP-BSA, or CPE in 100 μl of carbonate bicarbonate buffer (Sigma, C-3041), sealed with an adhesive cover, and incubated overnight in the fridge. The next day, plates were blotted entirely and blocked for 2 hours at room temperature (RT) with 100 μl of 5% skim milk powder or 1% BSA dissolved in PBS. Plates were washed 3 times with 0.05% Tween 20 in PBS (PBST) and incubated with 50 μl of serum diluted to the indicated concentrations in 1% skim milk or 0.3% BSA in PBS overnight in the fridge. The next day, plates were washed 3 times and incubated with 50 μl of either 0.25 μg/ml biotinylated anti-mouse IgG1 (Southern Biotech, 1070-08), or biotinylated anti-mouse IgG (H+L) (Poly4053, Biolegend, 405301) for 2 hours at RT. Plates were then washed 3 times and incubated with 50 μl streptavidin-alkaline phosphatase (ThermoFisher, 434322) diluted 1:1000 in 0.3% BSA-PBS for 1 hour at RT, covered from light. After 3 washes, plates were developed using 50 μl 4-nitrophenyl phosphate (Sigma, N- 9389) dissolved in 1x diethanolamine substrate buffer (ThermoFisher, 34064) and stopped using 25 μl of 1N NaOH. Plates were read at 405 nm.

#### IgE

Plates were coated with 50 μl of 2 μg/ml anti-mouse IgE (R35-72, BD, 553413) in PBS, sealed with an adhesive cover, and incubated overnight at 4℃ . The next day, plates were washed 3 times and blocked for 1 hour at 37℃ with 5% skim milk in PBS. Plates were washed 3 times and incubated with 50 μl of serum diluted to the indicated concentrations in 1% skim milk or 0.3% BSA in PBS overnight at 4℃. The next day, plates were washed 5 times, incubated with 50 μl of 300 ng/ml OVA, 150 ng/ml CPE, or 300 ng/ml NP-BSA conjugated to digoxigenin following supplier recommendations (ANP technologies, 90-1023-1KT) for 90 minutes at RT. Plates were washed 3 times, incubated with 50 μl of anti-digoxigenin POD fragments (Roche, 11 633 716 001) diluted 1:5000 in 0.3% BSA PBS for 1 hour at RT, covered from light. Plates were washed 5 times, then developed using 50 μl TMB (Sigma, T0440) and stopped using 25 μl 2N H2SO4. Plates were read at 450 nm.

#### Thiocyanate Elution

A thiocyanate elution step was added after sample incubation. Plates were washed 3 times, and wells were incubated for 15 minutes with the indicated concentrations of potassium thiocyanate (Sigma, 207799) at RT. Plates were then washed, and the ELISA protocol continued as described above.

### Degranulation Assay

Serum samples from allergic mice were diluted to 100 ng/ml based on total IgE. Total IgE was measured by ELISA as described above, however rather than coating the plate with antigen, the plate was coated with anti-IgE (LO-ME-3, Fitzgerald, 10R-I105A), and detection was performed using anti-IgE (23G3, Southern Biotech, 1130-08). Concentrations were determined using mouse IgE (C48-2, BD, 557080) as a standard. For heat-inactivation, serum was heated to 56°C for 1 hour.

Rat Basophil Leukemia cells (RBL-2H3, ATCC, Cat: CRL-2256) were incubated with diluted serum samples, plated in a 96 well flat bottom plate, and equilibrated at 37°C, 5% CO2 for 10 minutes. RBL-2H3 degranulation was assessed as previously described^51^. Briefly, sensitized RBL- 2H3 cells were stimulated with the indicated concentrations of allergen and incubated at 37°C for 30 minutes. Cell-free supernatant was collected, and cells were lysed with 0.1% Triton X-100. Cell supernatants and cell lysates were incubated in p-nitrophenyl N-acetyl-β-D-glucosamide in citrate buffer for 90 minutes at 37°C followed by the addition of 0.4 M glycine. Optical density at 405 nm was acquired using a Thermo Scientific Multiskan FC plate reader. Percent degranulation was calculated as 100 x (supernatant content)/(supernatant+lysate content).

### Tissue Collection and Processing

Mesenteric lymph nodes were collected into HBSS (Fisher, SH30015.03) and kept on ice until processing. Mesenteric lymph nodes were crushed between frosted microscope slides into a single cell suspension in HBSS, and passed through a 40 μm strainer (Corning Inc., 352340). After processing, single cells were pelleted by centrifugation and re-suspended in FACS buffer (2% FBS (Gibco, F4135), 2 mM EDTA (Sigma, 324504) in PBS) for downstream analysis. Cells were counted manually using a hemocytometer. Total cells were counted with Turks stain.

### Flow Cytometry

All steps were performed in FACS buffer unless otherwise specified. Three million cells were plated into a 96 well U-bottom plate (Corning, 353077) for staining. Centrifugation steps were performed at 200g at 4°C prior to fixation, and 300g after fixation. After each centrifugation the supernatant was decanted. Table 1 contains a list of all antibodies used for flow cytometry staining and the dilutions they were used at, based on 50 μl final staining volume.

**Table 1:** Antibody-fluorochrome conjugates and other reagents used for flow cytometry.

| Marker | Clone | Company | Conjugate | Dilution | Catalogue |
| --- | --- | --- | --- | --- | --- |
| B220 | RA3-6B2 | BioLegend | Alexa Fluor 700 | 100 | 103232 |
| CD138 | 281-2 | BioLegend | BV605 | 100 | 142516 |
| CD19 | 6D5 | BioLegend | PerCP-Cy5.5 | 100 | 115534 |
| CD95 | Jo2 | BD Biosciences | PE-Cy7 | 100 | 557653 |
| Fixable Viability Dye |  | ThermoFisher Scientific | eFluor 780 | 600 | 65-0865-18 |
| GL7 | GL7 | BioLegend | PerCP-Cy5.5 | 100-200 | 144610 |
|  |  | BioLegend | Alexa Fluor 647 | 100-200 | 144606 |
| IgD | 11-26c.2a | BioLegend | BV510 | 100 | 405723 |
| IgE | RME-1 | BioLegend | PE | 200-400 | 406908 |
| IgG1 | RMG1-1 | BioLegend | BV421 | 50 | 406616 |
| IgG2b | RMG2b-1 | BioLegend | Alexa Fluor 488 | 100 | 406718 |
| IgG2c |  | Southern Biotech | FITC | 100 | 1079-02 |
| IgG3 | R40-82 | BD Biosciences | BV650 | 100 | 744136 |
| IgM | II/41 | BD Biosciences | BV786 | 50 | 743328 |
| OVA |  | ThermoFisher Scientific | FITC | 100 | O23020 |
|  |  | ThermoFisher Scientific | Alexa Fluor 647 | 100 | O34784 |

Cells were pelleted and resuspended in 25 μl FACS with anti-CD16/32 (Clone 93, Biolegend, 101302), to block antibodies binding to Fcγ receptors, for 15 minutes on ice. In experiments where intracellular IgE was detected, unlabeled anti-mouse IgE (Clone RME-1, Biolegend, 406902) was used to block surface IgE bound to the low affinity IgE receptor (CD23) expressed by B cells. Twenty-five μl of a mix of antibodies to label extracellular targets was added directly to blocking solution, and the cells were incubated for 30 minutes on ice, covered from light. After this incubation, cells were washed twice by adding 200 μl of FACS and centrifuging each time. For fixation and permeabilization, cells were resuspended and incubated in 100 μl of BD Cytofix/Cytoperm (BD, 554714) for 20 minutes. Cells were then washed with 150 μl BD Perm/Wash (BD, 554714), pelleted, washed again with 200 μl BD Perm/Wash, and pelleted again. For intracellular staining, cells were then resuspended in 50 μl of intracellular staining cocktail in BD Perm/Wash and stained for 30-45 minutes on ice, protected from light. Finally, cells were washed twice with 200 μl of BD Perm/Wash and then resuspended in 200 μl of FACS. Data were collected using a BD LSRFortessa (BD, Franklin Lakes, USA), and analyzed using FlowJo (FlowJo LLC, Ashland, USA). Allergen-specific cells were probed using FITC-conjugated OVA (Molecular Probes, O23020) in the extracellular staining mix.

### Statistical Analysis

Experiments were performed with multiple biological replicates and multiple times (see figure legends) to ensure reproducibility and statistical power to detect meaningful differences. GraphPad Prism v9 was used for all statistical analyses. Data which approximated a log-normal distribution were log transformed prior to statistical testing, except for data sets which included values of 0 (*e.g.*, clinical sign data) which could not be log-transformed. See figure legends for specific statistical tests performed.

## Acknowledgements

We thank Hong Liang from the McMaster Flow Cytometry Core for assistance with flow cytometry experiments. We thank McMaster’s Central Animal Facility for support with animal experimentation. We thank Amy L. Kenter (University of Illinois) for generously providing hMT mice. We thank Dr. Hirohito Kita (Mayo Clinic) for critical review of the manuscript prior to publication.

## Funding

Schroeder Foundation (JFEK, SW, MJ) Food Allergy Canada (JFEK, SW, MJ) ALK Abello A/S (JFEK, MJ)

Zych Family (SW, MJ) Satov Family (SW, MJ)

Canadian Allergy Asthma and Immunology Foundation (MJ) Canadian Society of Allergy and Clinical Immunology (MJ) Canadian Institutes of Health Research (JFEK)

Canadian Institutes of Health Research Doctoral Foreign Study Award DFD-17076 (AKWV)

European Social Fund and Instituto de Salud Carlos III through a Miguel Servet grant CP20/00043 (RJS)

Canadian Institutes of Health Research Doctoral Research Award & Fellowship (KB)

## Author Contributions

Conceptualization: JFEK, AKWV, RJS, MJ

Formal Analysis: JFEK, AKWV, KB

Investigation: JFEK, AKWV, RJS, KB, SG, TW, MG

Visualization: JFEK, AKWV

Funding Acquisition: JFEK, SW, MJ

Supervision: JFEK, SW, MJ

Writing – Original Draft: JFEK, AKWV

Writing – Review & Editing: JFEK, AKWV, RJS, MJ

---

**Supplementary Figure 1:**
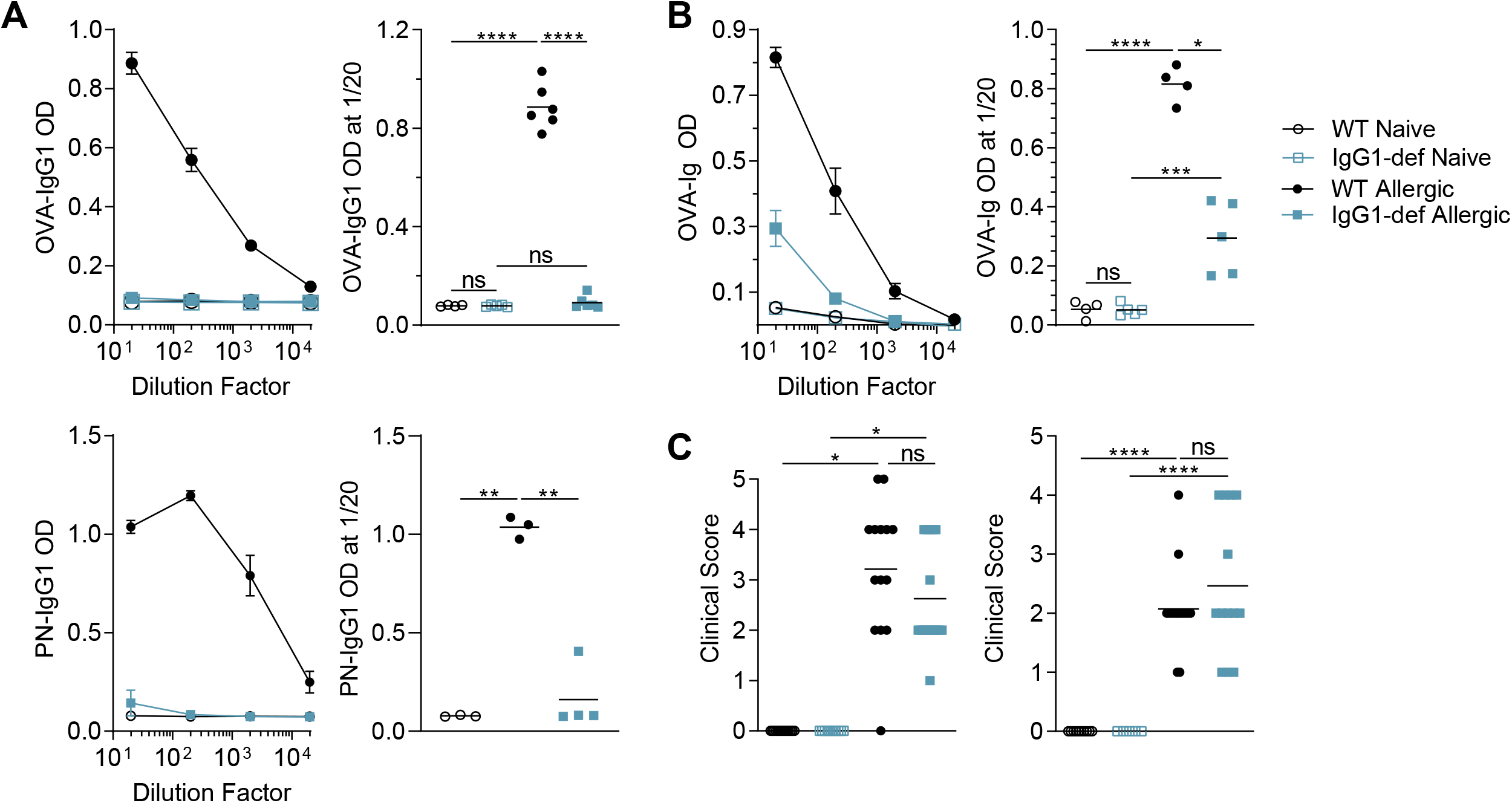
Supporting information for Figure 1. (A-C) Sensitization was performed as described in Figure 1 and Materials and Methods. **(A)** Serum dilution curve for OVA- specific (top row) or PN-specific (bottom row) IgG1 assessed by ELISA at 450nm (left column) with statistical analysis at the 1/20 dilution (right column). **(B)** Serum dilution curve for total OVA- specific Ig (left) with statistical analysis at the 1/20 dilution (right). **(C)** Assessment of clinical symptoms of anaphylaxis in the 40 minutes following challenge with allergen in experiments using OVA (left) or PN (right). **(D)** Clinical scores for signs of anaphylaxis upon challenge; same across all figures. *, P<0.05; **, P<0.01; ***, P < 0.001; ****, P < 0.0001 (one way ANOVA with Tukey’s post-test comparing the mean of each group with the mean of every other group [A-C]). Statistical notations atop bars indicate comparisons between the bridged groups. Dots represent group means (A – left, B – left), measurements of samples from individual mice (A – right, B – right), or readings from individual mice (C). Bars within groups represent the mean. Data are representative (A-B) or pooled from (C) three independent experiments.

**Supplementary Figure 2:**
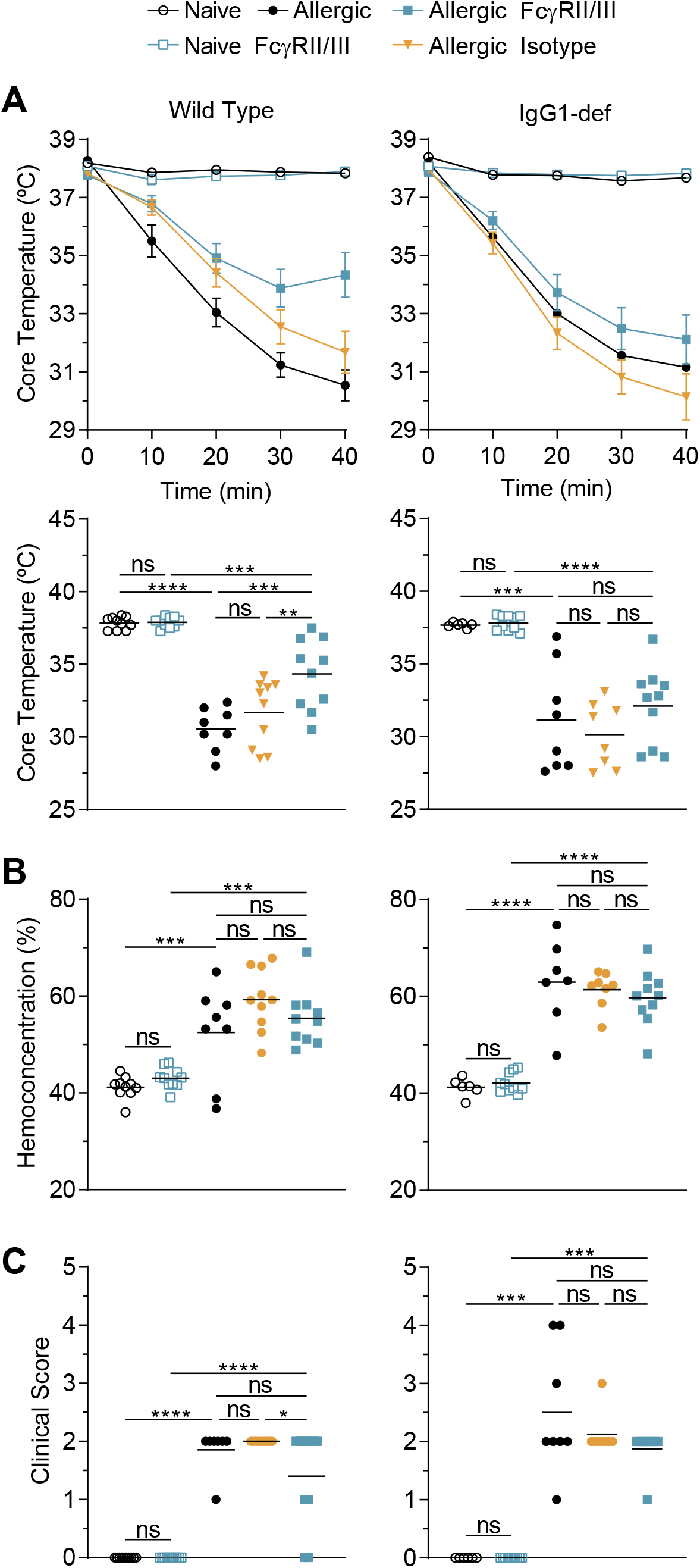
Anaphylactic symptoms of IgG1-def mice are mediated by IgE. (A-C) Naïve and PN-sensitized mice were left untreated or were treated with anti-FcγRII/III or isotype control 1 d prior to challenge. Results for WT and IgG1-def mice are depicted in the left and right columns, respectively. **(A)** Measurement of core temperature change over time following intraperitoneal allergen challenge (top row) with statistical analysis at 40 minutes post-challenge (bottom row). **(B)** Quantification of hematocrit in blood collected at 40 minutes post-challenge. **(C)** Assessment of clinical symptoms of anaphylaxis in the 40 minutes following allergen challenge. *, P<0.05; **, P<0.01; ***, P < 0.001; ****, P < 0.0001 (one way ANOVA with Tukey’s post-test comparing the mean of each group with the mean of every other group). Statistical notations atop bars indicate comparisons between the bridged groups. Dots represent group means (A – top row), readings from individual mice (A – bottom row, C), or measurements of samples from individual mice (B). Bars within groups represent the mean. Data are pooled from two independent experiments.

**Supplementary Figure 3:**
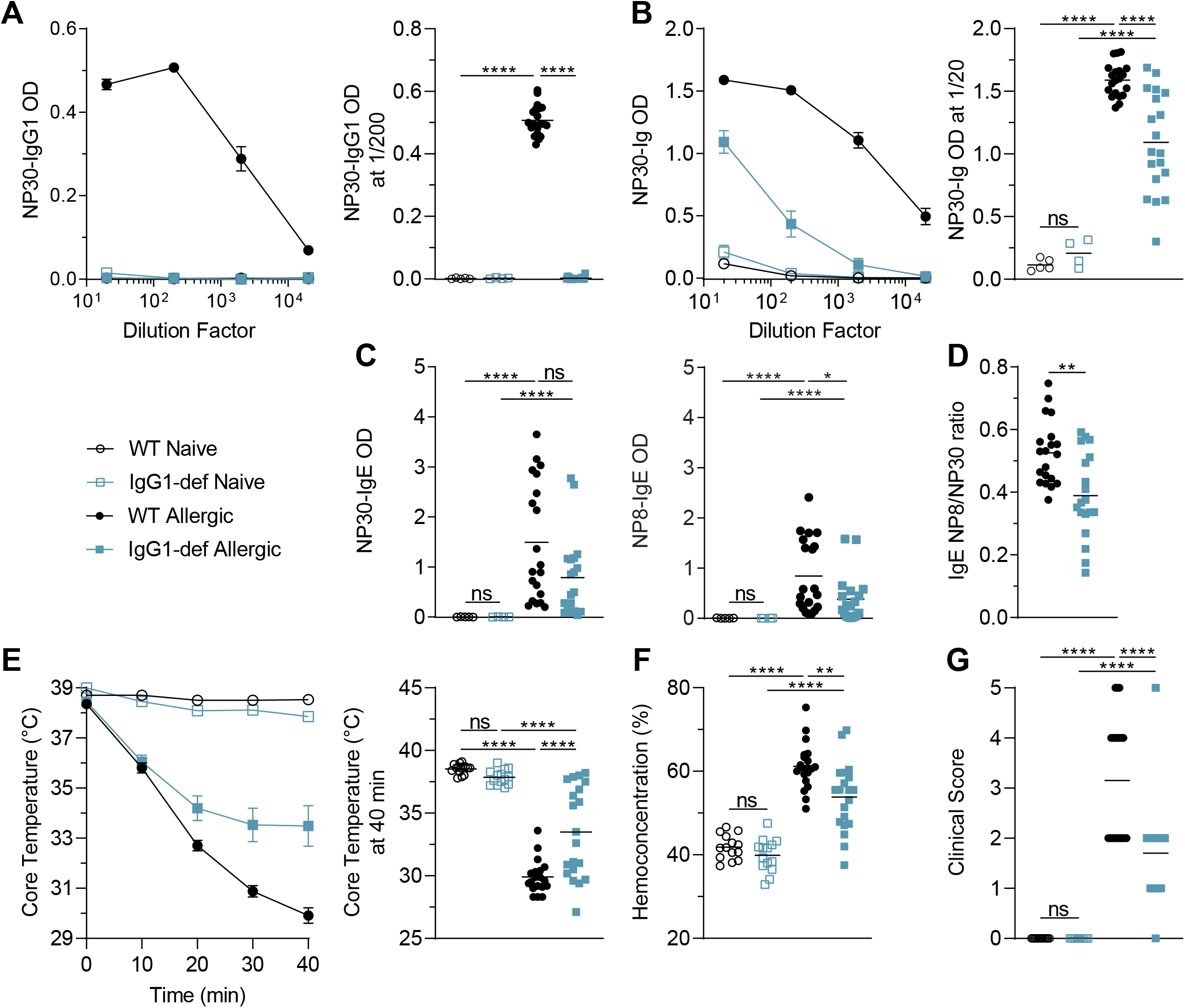
IgG1-def mice are partially protected from allergic reactivity against the hapten NP. (A-G) IgG1-def and WT mice were sensitized to NP-OVA. **(A-C)** Serum dilution curves (left plots) for NP(30)BSA-specific IgG1 (A), total NP(30)BSA-specific Ig (B), or NP(30)BSA-specific IgE (C) assessed by ELISA at 450nm (A-B) or 405nm (C) with statistical analysis (right plots) at the 1/20 dilution (A-B) or 1/2 dilution (C). **(D)** The ratio of the OD of an NP(8)BSA-specific IgE ELISA divided by the OD of an NP(30)BSA-specific IgE ELISA per sample. **(E)** Core body temperature over time following challenge with NP(30)BSA (left) with statistical analysis at the 40-minute timepoint (right). **(F)** Evaluation of hematocrit in blood collected at 40-minutes post-challenge. **(G)** Grading of clinical signs of anaphylaxis in the 40 minutes following allergen challenge. **, P<0.01; ****, P < 0.0001 (one way ANOVA with Tukey’s post-test comparing the mean of each group with the mean of every other group [A-C, E-G], unpaired t test [D]). Statistical notations atop bars indicate comparisons between the bridged groups. Dots represent group means (A – left, B – left, E – left), or readings from individual mice (A – right, B – right, C-D, E – right, F-G). Bars within groups represent the mean. Data are pooled from three independent experiments.

